# Joint testing of rare variant burden scores using non-negative least squares

**DOI:** 10.1101/2023.02.22.529560

**Authors:** Andrey Ziyatdinov, Joelle Mbatchou, Anthony Marcketta, Joshua Backman, Sheila Gaynor, Yuxin Zou, Tyler Joseph, Benjamin Geraghty, Joseph Herman, Kyoko Watanabe, Arkopravo Ghosh, Jack Kosmicki, Adam Locke, Regeneron Genetics Center, Timothy Thornton, Hyun Min Kang, Manuel Ferreira, Aris Baras, Goncalo Abecasis, Jonathan Marchini

## Abstract

Gene-based burden tests are a popular and powerful approach for analysis of exome-wide association studies. These approaches combine sets of variants within a gene into a single burden score that is then tested for association. Typically, a range of burden scores are calculated and tested across a range of annotation classes and frequency bins. Correlation between these tests can complicate the multiple testing correction and hamper interpretation of the results. We introduce a new method called the Sparse Burden Association Test (SBAT) that tests the joint set of burden scores under the assumption that causal burden scores act in the same effect direction. The method simultaneously assesses the significance of the model fit and selects the set of burden scores that best explain the association at the same time. Using simulated data, we show that the method is well calibrated and highlight some scenarios where the test outperforms existing gene-based tests. We apply the method to 73 quantitative traits from the UK Biobank which further illustrates the power of the method. This test is implemented in the REGENIE software.

## Introduction

Large scale exome sequencing studies are being conducted to elucidate the genetic basis of diseases and traits and discover novel drug targets^1–3^. These studies enable association testing of rare coding variation not easily accessible via genome-wide association studies (GWAS) using genotype microarrays followed by imputation from publicly available reference panels. Typically, these studies will carry out statistical tests of the combined effect across many single variants in each gene with a trait of interest.

Possibly the simplest approach involves collapsing a subset of single variants in a gene into a single marker of gene perturbation or burden^4^. For example, the set of predicted loss of function (pLoF) variants below 0.1% minor allele frequency (MAF) could be combined by scoring an individual as 1 if they have at least one minor allele across the set, and 0 otherwise. This *burden score* can then be tested for association in the same way as a SNP (single nucleotide polymorphisms). Alternatives include a weighted sum of variants, with weights dependent upon the MAF of variants^5^, or a set of burden scores across a range of frequency thresholds, with significance assigned using permutation^6^.

Burden tests tend to have most power when all collapsed variants are causal and when causal variants alter gene function in the same effect direction. When these conditions are not met variance components tests, such as the SKAT test^7^, that allow variants to have different effect directions can have more power. Alternatively, combining single variant tests, via the Cauchy combination test^8^ (called ACAT-V), into a single p-value can be particularly powerful when there are only a small number of causal variants ^9^. Tests that combine across burden, variance component and single variant tests have also been proposed and exhibit good power^9–11^.

As the true set of causal variants is unknown it is common to calculate many burden scores across a range of MAF thresholds and different variant annotation classes^2,12^. For example, Backman et al. ^2^ considered two annotation classes and five MAF thresholds: a strict burden of pLoFs and a more permissive burden of pLoFs with predicted deleterious missense variants were assigned into overlapping groups with MAF ≤ 1%, MAF ≤ 0.1%, MAF ≤ 0.01%, MAF ≤ 0.001%, and singletons only. This approach tends to produce highly structured, and often highly correlated sets of tests, which can make the interpretation and accurate multiple testing correction difficult.

In this paper we propose an approach that circumvents these difficulties whereby a set of burden scores are *jointly tested* for association. We focus on quantitative traits and leave the extension to binary traits for future work. As the set of nested burden scores tend to be positively correlated and exhibit the same direction of association with the trait of interest this suggests a prior on the effect direction of the burden scores, which we enforce through fitting the joint model of burden scores using a non-negative least squares (NNLS) approach. We propose a quadratic form test statistic based on the NNLS model fit and show that it has a null distribution that is a *mixture* of chi-square distributions (not to be confused with a *weighted sum* of chi-squares) with mixture weights that depend on the covariance between the burden scores. We also develop a computationally efficient method for calculating p-values from this null distribution. NNLS is known to induce sparsity in its solution and has the added benefit of providing a form of model selection across the set of burden scores, which can aid interpretation of which frequency bins and annotation classes harbor the causal variants. In this sense the method is rather unique in simultaneously achieving both model inference, via well controlled Type I error for the test, and model selection, with many of the burden scores excluded from the final model fit. We call this test the Sparse Burden Association Test (SBAT).

Using simulation studies, we show that the SBAT has well calibrated Type I error and highlight some scenarios where the SBAT has improved power over existing tests. We further show the performance of the test when applied to 73 quantitative traits from the UK Biobank and illustrate how the sparse nature of the parameter estimation in the test can aid interpretation of the causal signal at a gene.

## Methods

### Sparse Burden Association Test

In a sample of *N* individuals, let *Y* denote the *N*-length vector of a quantitative trait. Let *G* denote the *N* × *L* matrix, where each column contains the genotypes of *L* SNPs in a gene of interest. Further define *X* to be a matrix of *N* × *P* burden scores, where each column contains one of *P* distinct burden scores. Each burden score is derived as function of the SNP matrix *X*, by collapsing the subset of variants that fall within a MAF bin and an annotation class. That is *X*_*ij*_ is the indicator variable that the *i*th individual carries at least one rare variant at any of the SNPs included in the *j*th subset. The set of MAF bins and annotation classes are user defined. We assume that covariates (including the intercept) are projected out from both *Y* and *X*. Covariates may include terms such as REGENIE Step 1 predictors^13^ that are estimated in advance and account for polygenicity, population structure and relatedness.

We consider estimating the joint effect of all *P* burden scores in a linear model with positivity constraints on the *P*-vector of regression parameters *β* as follows

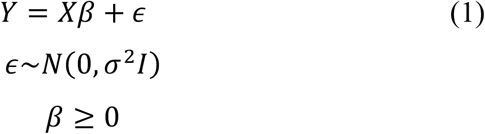

We assume that *σ*^2^ is known or is estimated with enough precision so that this can be ignored, such as the case of large-scale exome sequencing datasets. This is an NNLS problem and we use the active set method^14^ to fit the model. We use 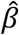 to denote the NNLS estimate and 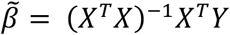 to denote the ordinary least squares (OLS) estimate of *β* without the non-negativity constraint.

Writing 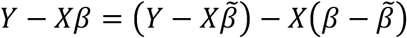 the least squares objective function for (1) can be re-expressed as

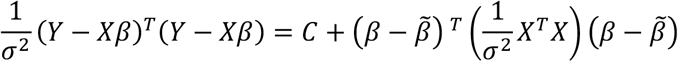

Where 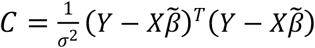 and cross product terms vanish as a consequence of the OLS normal equation 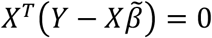. Hence model (1) is equivalent to

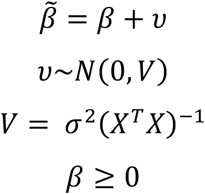

Treatment of this model is explicitly addressed by literature on inference of linear models with constraints^15–17^, and the model can be thought as a multivariate analogue of the one-sided test ^15^. To test the null hypothesis *H*_0_: *β* = 0 versus the alternative *H*_1_: *β* ≥ 0 (with at least one inequality strict) we use the quadratic form test statistic

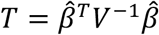

That has a null distribution that is a *mixture* of chi-squared distributions^15–17^ (not to be confused with a *weighted sum* of chi-squares) as follows

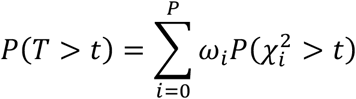

Where 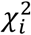 is a chi-squared distribution with *i* degrees of freedom, and *ω*_*i*_ are weights that sum to 1 and depend upon *V* as follows

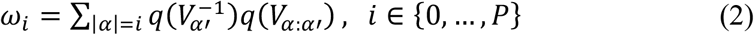

over all subsets of {1, …, *P*} of size *i* denoted by *α*, and *α*′ is the complement set. *V*_*α*′_ is the covariance matrix of *υ*_*i*_, *i* ∈ *α*′ and *V*_*α*:*α*′_ is the covariance matrix of *υ*_*i*_, *i* ∈ *α* conditional upon *υ*_*i*_ = 0, *i* ∈ *α*^′^, and *q*(Σ) is the probability that all the variables of a multivariate normal distribution with zero mean and covariance Σ are positive. In the case where *V* is diagonal the weights simplify to

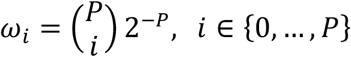

In the general case the sum *ω* has 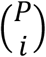 terms to evaluate which can be a potential computational bottleneck for P > 10. For example, *ω* for P = 25 implies evaluating 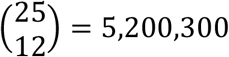 terms in the sum from Equation 2. Thus, we explored a simple approximation in which a random sample of *K* terms in each sum, denoted *A*_*i*_, were used to estimate the full sum

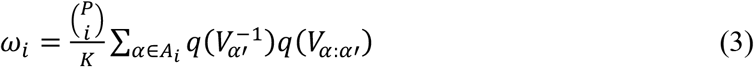

As we do not know in advance whether the burden scores will be positively or negatively associated with the quantitative trait under study, we apply the SBAT twice to both *Y* and −*Y*. We then combine the two p-values using the Cauchy combination method^8^. To avoid numerical instability, we apply the QR decomposition to the matrix of burden scores and leave only linearly independent columns for model fitting and inference.

This approach, together with a general suite of burden, variance component and ACAT (Aggregated Cauchy Association Test) gene-based tests have been implemented in the REGENIE version 3.0 software (**URL**s), and this was used for all simulation and real data analysis results included in this paper.

### Annotations and MAF cutoffs for burden scores

In both simulations and analysis of the UK Biobank data, exome variants were grouped based on four annotations classes: (1) pLoF only (labeled M1); (2) pLoF or predicted deleterious missense based on 5/5 in-silico algorithms (labeled M3); (3) pLoF or predicted deleterious missense based on 1 or more out of 5 in-silico algorithms (labeled M4); (4) and pLoF or missense (labeled M2). We also considered several MAF cutoffs when aggregating variants: MAF ≤ 1%, MAF ≤ 0.1%, MAF ≤ 0.01%, MAF ≤ 0.001% (only for UK Biobank application), and singletons. Exome variants were annotated using SnpEff and assigned to genes based on Ensembl v85 (most deleterious consequence across any transcript) as previously described^2^.

### Combining different gene-based tests into a single p-value per gene

We considered four gene-based tests: SBAT, SKAT-O^10^ and ACAT-V^9^, and another test, BURDEN-ACAT, that aggregates evidence of association from multiple burden scores using the Cauchy combination method with uniform weights^8,9^. For the SBAT and BURDEN-ACAT tests, input was a set of burden scores built by grouping exome variants using four annotation classes and five MAF cutoffs as described above. For the SKAT-O and ACAT-V tests, we applied a single MAF cutoff of 1% along with MAF-dependent variant weights and aggregated the resulting p-values (across the four annotation classes) by the Cauchy combination method. That produced a single per-gene p-value for each of the four gene-based tests. Finally, we applied the Cauchy combination method on the four p-values to derive a single p-value per gene, referred to as GENE_P (**Supplementary Figure S3**).

### Simulated data

To assess the Type I error calibration of the SBAT, we simulated phenotypes based on real genetic data from the UK Biobank array^18^ and exome data^2^ as follows. We randomly selected 100,000 samples from the set of white British participants to include population structure in our simulations (about 30% of samples are related 3^rd^ degree or closer). For the *i*-th individual, quantitative phenotypes were simulated as

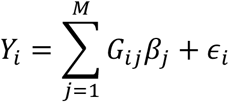

where we selected *M=*25,000 SNPs on odd chromosomes from the UK Biobank genotyping array to be causal, excluding variants with minor allele count below 100 or involved in inter-chromosomal LD (Linkage Disequilibrium), and *G*_*ij*_ is the standardized genotype value at the *j*-th marker for the *i*-th individual. The effects for the causal SNPs *β*_*j*_ were sampled independently from a normal distribution with mean zero and the variance was chosen so that they explained 20% of the total phenotypic variance. The environmental effect *ϵ*_*i*_ was independently sampled from a normal distribution with mean zero and variance set to result in a phenotypic variance of one. We simulated 100 phenotypic replicates, obtained REGENIE Step 1 predictors based on 472,435 array SNPs, and tested the phenotypes for association with 1,000 randomly selected genes on even chromosomes from whole exome sequencing data to evaluate type 1 error.

We also used UK Biobank array and exome data as the basis for a set of simulations to assess the power of the SBAT compared to SKAT-O^10^ and ACAT-V^9^ tests. For the *i*-th individual, quantitative phenotypes were simulated as

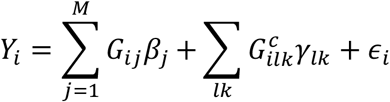

where we followed the same scheme as in the type 1 error simulations to simulate additive polygenic effects from array SNPs on odd chromosomes, but also added effects from causal genes selected on even chromosomes, where 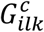 is the genotype value (non-standardized) for the *l-*th exome variant in the *k*-th causal gene for the *i*-th individual, and *γ*_*lk*_ the corresponding fixed effect on the phenotype. For each causal gene, we only considered variants that were annotated either pLoF or missense. The absolute effect sizes |*γ*_*lk*_| were |0.1log_10_(MAF)| for pLoF variants, and |0.01log_10_(MAF)| for missense variants. This reflected the assumption that functional variants which are more deleterious are likely have larger effect sizes on the phenotype and be rarer. For singleton variants, we set the effect size to *γ*_*lk*_*R*_*singleton*_, where *R*_*singleton*_ ≥ 1 is a positive constant which enabled singletons to have more severe impact on the phenotype than that solely based on the MAF. We varied the proportion of causal variants among non-singleton variants between 10 to 100% and for singleton variants between 30 to 100%. For the direction of effects, we considered three settings where (1) all variants have positive effects; (2) 80% of variants have positive effects (and remaining 20% had effects in the opposite direction); and (3) 50% of variants have positive effects. In all scenarios, the polygenic effect from array SNPs was set to explain 20% of the phenotypic variance when there are no effects from the causal genes (i.e., all *γ*_*lk*_ = 0) with the remaining 80% of the variance explained by the environmental effect. We simulated 100 phenotypic replicates which we tested for association with each of the 10 causal genes resulting in 1,000 p-values and evaluated power in each scenario. Similarly to the type 1 error simulations, we also obtained REGENIE Step 1 predictors based on 472,435 array SNPs before scanning for associations.

### Analysis of UK Biobank data

We applied SBAT to real data analysis of 73 quantitative phenotypes in the UK Biobank with sample sizes ranging from 89,734 to 430,074 European participants and up to 18,184 genes on autosomal chromosomes; these included biomarkers as well as anthropometry outcomes (**Supplementary Table S1**). These phenotypes were derived from the phenotypes available through the UK Biobank Data Showcase on April 1, 2020. For traits measured across multiple visits, we computed the mean value across all visits for each participant and analyzed the resulting variable after applying a rank-based inverse-normal transformation. For each trait, we first performed a GWAS with imputed variants to identify common variants independently associated with the phenotype as described in Backman et al^2^. Briefly, we tested common (MAF≥1%) single variants imputed from TOPMed (Trans Omics for Precision Medicine) for association with the trait using REGENIE. We then ran GCTA-COJO^28^ joint model to identify independent signals (*P* ≤ 10^−7^) using 10,000 randomly selected individuals from the UK Biobank TOPMed dataset to estimate linkage disequilibrium. These independently associated variants were included as covariates in the set-based test analyses we performed on exome rare variants. For the gene-based association tests, we grouped variants into 20 burden scores using the same four annotation classes as described above and the MAF cutoffs of 1%, 0.1%, 0.01%, 0.001% and singletons. For SKAT-O and ACAT-V association tests, we only considered a single MAF cutoff of 1%. Covariates included age, age-squared, sex, age-x-sex, the first ten principal components based on array data, the first twenty principal components derived from exome variants with a MAF between 2.6 × 10^−5^ and 1%, and six exome sequence batch indicator variables.

## Results

### Approximate SBAT p-values

To mitigate the computational burden of weight calculation for SBAT (Equation 2), we propose approximation with a single parameter K that controls the size of the random sample of terms in the weight calculation (Equation 3). We evaluate the performance of our approximation using three values of K = 1, 10, 100 (**Figure 1**). The accuracy of weight approximation naturally improves with K increasing from 1 to 100, as highlighted for 12 burden scores of the *APOB* gene in the UK Biobank (**Figure 1a**). We next expand the comparison to 812 genes on Chromosome 21 in the UK Biobank tested for association with low-density lipoprotein (LDL) phenotype (**Figure 1b-c**). The exact and approximate p-values are almost undistinguishable when K is large enough (R^2^ >0.9999 for K=100), but differences between p-values are noticeable at very small values of K (R^2^ = 0.990 for K=1). We recommend the value of K = 10 (default in REGENIE), a reasonable compromise between the accuracy of approximation and compute time. To further reduce the computational burden of SBAT, we develop an adaptive strategy, named SBAT-ADAPT, which initially uses K=2 weights to estimate the p-value, and if it is below a significance level of 0.001, we re-compute the p-value using K=10 weights. This strategy allows to quickly evaluate p-values for weaker signals and increase accuracy for stronger signals which we are more interested in. **Supplementary Figure 1** shows the high concordance in p-values between this adaptive strategy and using K=10 weights to calculate all the p-values (R^2^ = 0.991). Furthermore, it led to a 40% reduction in computational time when performing a whole-exome association scan on a single phenotype compared to using K=10 weights (**Supplementary Table S3**). As REGENIE can analyze multiple phenotypes within the same analysis run^13^, we propose another strategy, named SBAT-MTW, which is better suited for this context. More precisely, instead of evaluating K weights separately for each trait, we compute the weights for the first phenotype and re-use these across all phenotypes. Assuming that the missingness patterns across traits is similar (which is also assumed in the REGENIE multi-trait setting), the LD matrices for the burden scores obtained given each phenotype’s missingness patterns should be highly correlated and thus lead to similar weights. We evaluate this strategy in real-data applications where SBAT-MTW is applied to 50 phenotypes and gives p-values highly concordant with those from SBAT (R^2^ = 0.997). **Supplementary Table S3** shows the timing reduction of SBAT-MTW relative to SBAT when applied to multiple phenotypes where it increased as more phenotypes are being analyzed in the same REGENIE run (CPU time reduction of 65% for the multi-trait run with 50 phenotypes).

**Figure 1.**
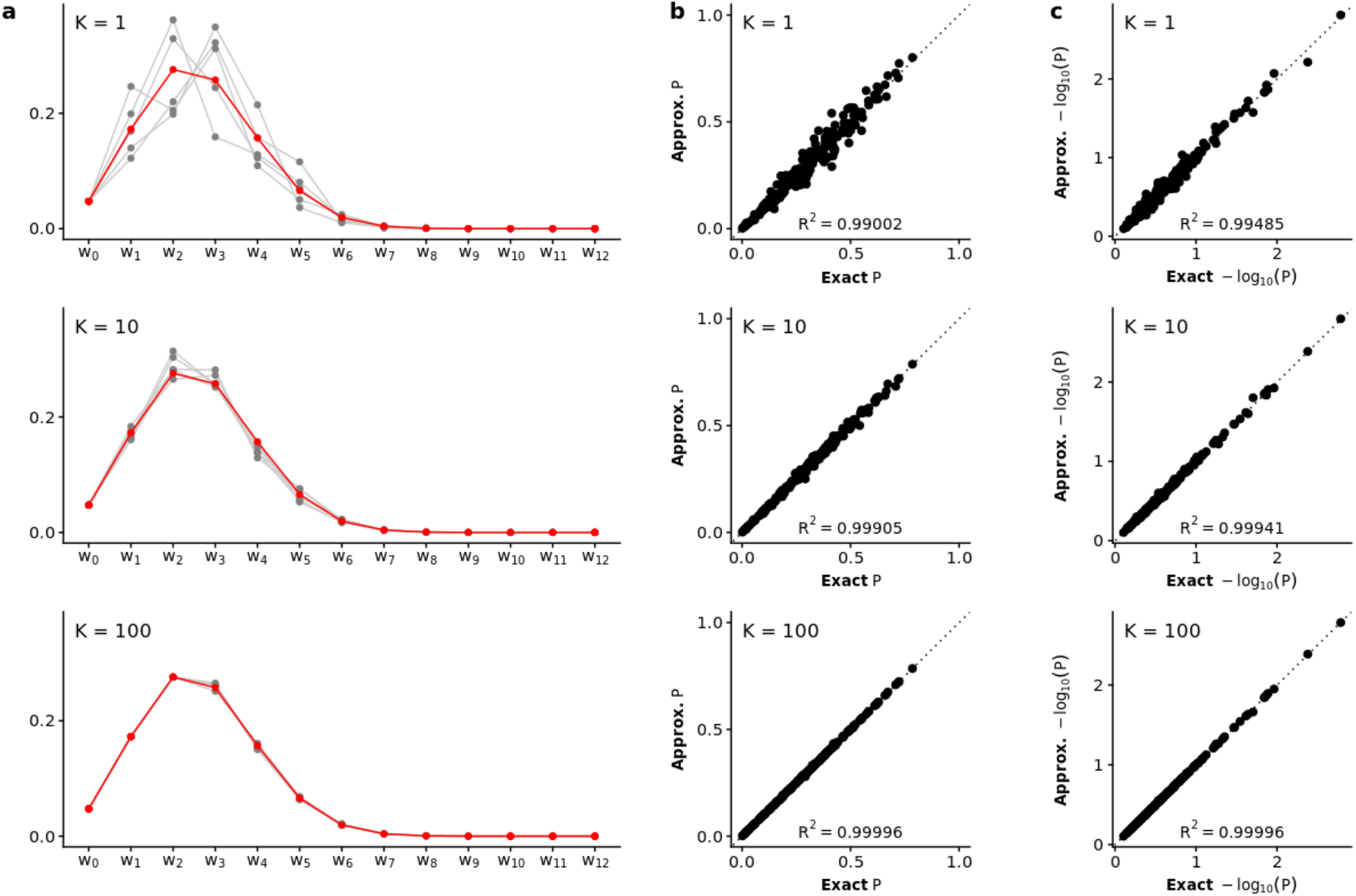
Validation of approximate SBAT p-values with different values of parameter K = 1, 10, and 100. (a) Exact (red) and approximate (grey) weights for 12 burden scores of the *APOB* gene in the UK Biobank are compared with 5 repetitions (grey curves). (b-c) Scatter plots of exact and approximate SBAT p-values at raw and log10 scale for 812 genes on Chromosome 21 and LDL phenotype in the UK Biobank. The R^2^ values are reported for both raw and log10-transformed p-values.

### Type 1 Error Simulation for SBAT

The QQ-plot on the log scale comparing the SBAT p-values (using *K* = 10) to the expected values under the null hypothesis of no association based on 100,000 null simulations is shown on **Figure 2**. The empirical type 1 error rates for SBAT for significance levels *α* = 0.05, 0.01, 0.005, 0.001 and 5 × 10^−4^ are presented in **Supplementary Table S2**. These results confirmed that the SBAT has correct type 1 error control for quantitative traits.

**Figure 2.**
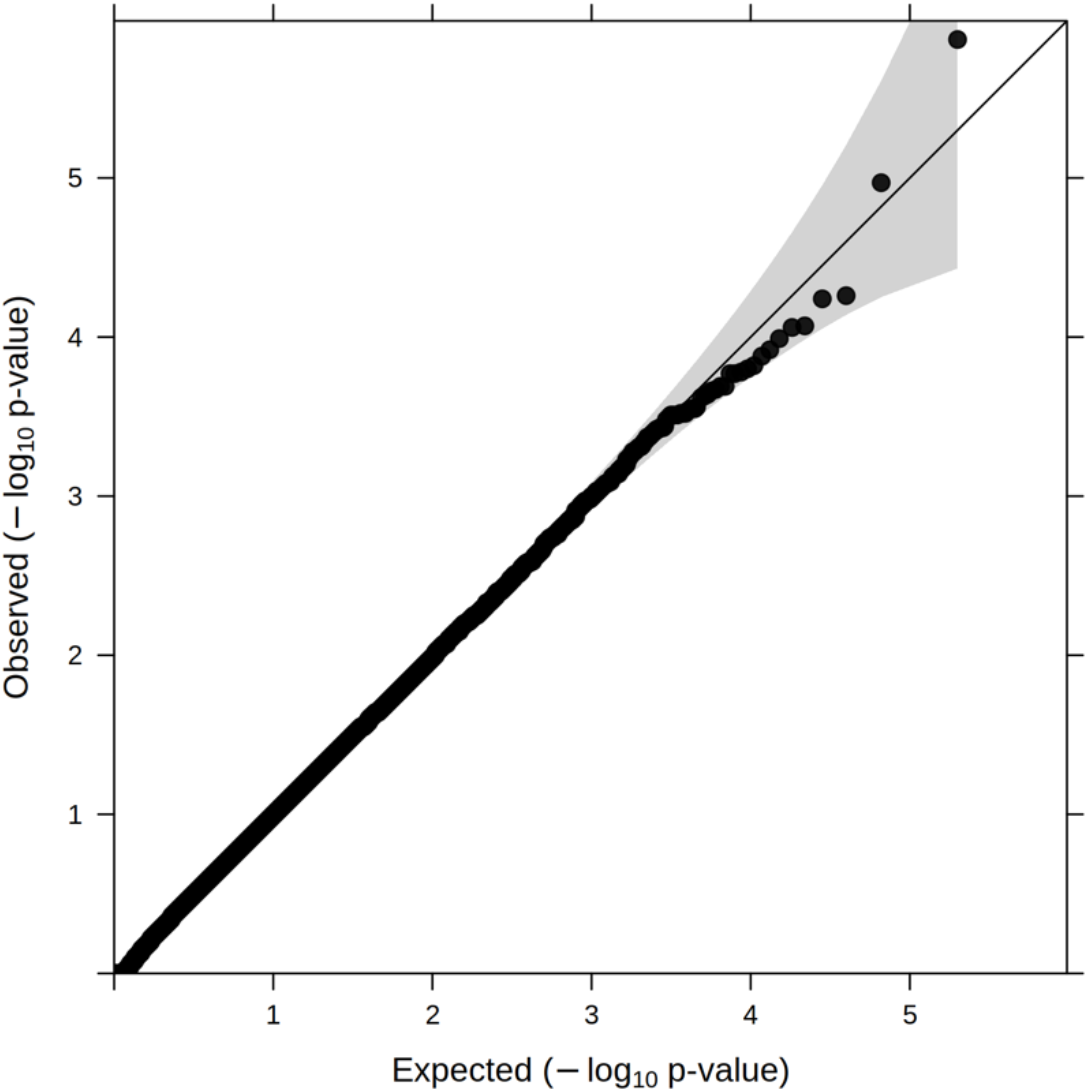
Quantile-quantile plot of association test p-values for SBAT in simulation studies under the null hypothesis. The p-values were obtained from testing 100 simulated phenotypes in 1,000 genes where variants were grouped by four functional annotations and four MAF cutoffs resulting in 16 burden scores being combined in SBAT.

### Power Simulation

We compared the power performance of SBAT with SKAT-O and ACAT-V under multiple simulation settings where we varied the proportion as well as the direction of causal signals based on their functional annotation. **Figure 3** shows the empirical power performance of the tests across a range of 6 simulation configurations. The SBAT test had the highest power relative to SKAT-O and ACAT-V when the burden assumption of same direction of effects for all the variants in the gene was held and singleton variants had significantly higher impact on the phenotype than non-singleton variants. However, the power performance of SBAT was diminished when only a fraction of the singleton variants was causal. We also found that when the increased effect of singleton variants was present, SBAT remained as powerful or more than SKAT-O and ACAT-V even when variant effects were not all in the same direction (80/20 +/-). We found SBAT had the lowest power performance in the settings that were the farthest from the burden assumption (50/50 +/-). In summary, SBAT performed better than SKAT-O and ACAT-V when the same effect direction assumption was met, and rarer variants were driving most of the gene signal.

**Figure 3.**
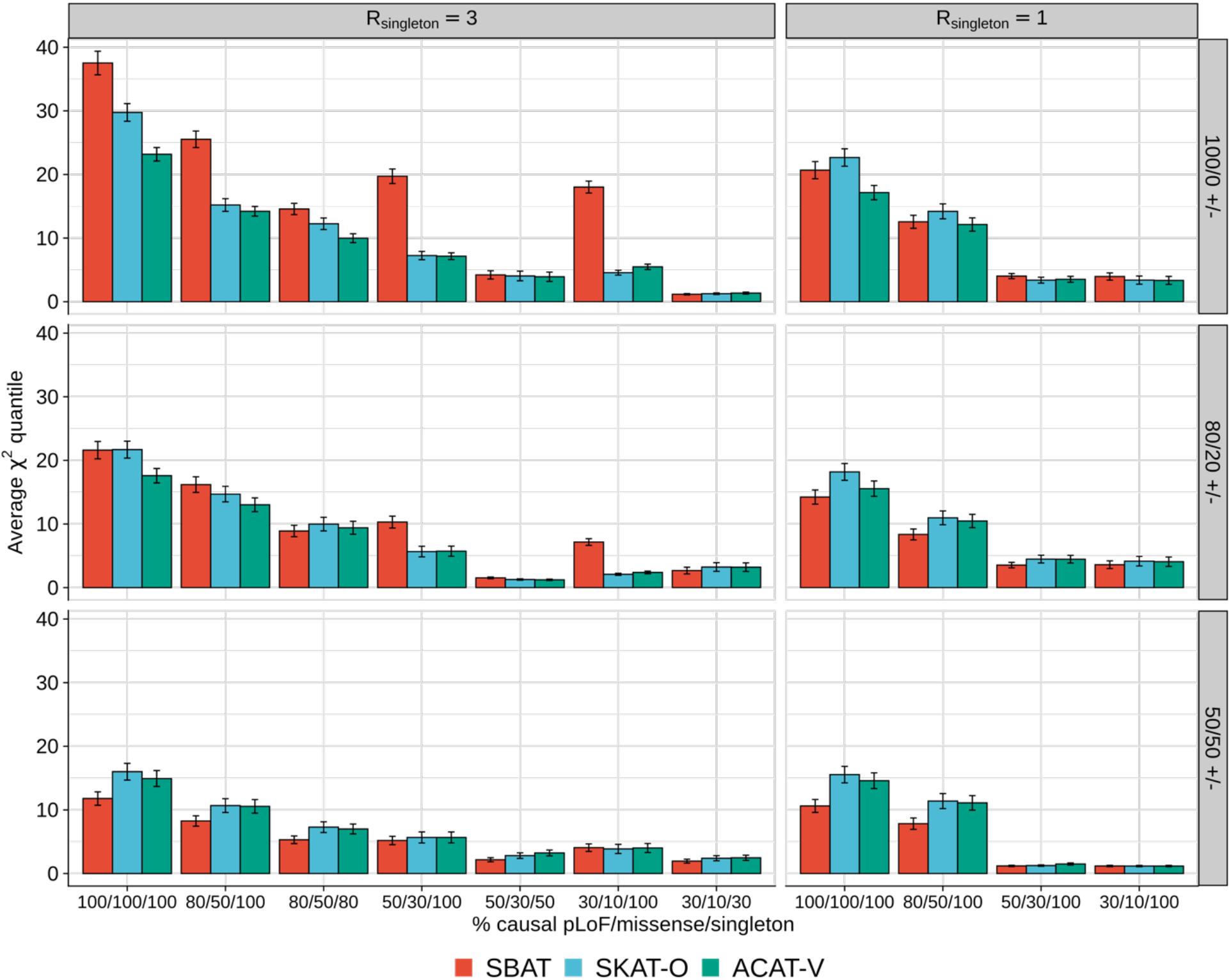
Power performance of SBAT in simulation studies under various causal scenarios. Average *χ*^2^ quantile for the set-based tests as a function of the proportion of causal variants amongst singletons, pLoF and missense variants. Each row corresponds to different assumptions for the direction of effects whether they are all positive (100/0 +/-), 80% positive and 20% negative (80/20 +/-), or half positive and the remaining negative (50/50 +/-). Each column corresponds to different assumptions for the effect of singleton variants, which was multiplied by a constant factor *R*_*singleton*_ relative to non-singletons. The error bars represent 95% confidence intervals for the average *χ*^2^ quantile.

### Application to UK Biobank

We analyzed whole-exome sequencing data on up to 18,184 genes for 73 quantitative phenotypes in up to 430,074 European participants from the UK Biobank using SBAT, BURDEN-ACAT, SKAT-O and ACAT-V association tests and conditioning on common variant signals identified through GCTA-COJO. We adjusted for age, sex, exome sequence batch variables as well as principal components derived from array and rare exome variants. We used a conservative genome-wide significance threshold of *α* = 9.4 × 10^−9^ accounting for the 73 quantitative traits, 18,184 genes and 4 association tests applied. The median runtime per trait was 279 CPU hours to analyze all 18,184 genes with SKAT-O, ACAT-V, BURDEN-ACAT and SBAT on 20 burden scores conditioning on an average of 644 common variants (the highest compute timing was for sitting height at 538 CPU hours with 1,722 conditional variants).

Across the 73 traits, SBAT identified 1,339 genome-wide significant associations, compared to 1,274 for BURDEN-ACAT, 1,462 for SKAT-O and 1,175 for ACAT-V. SBAT uniquely identified 115 signals, which was greater than the signals uniquely identified by BURDEN-ACAT and SKAT-O, 16 and 58, respectively, and lower than ACAT-V which had 158 unique signals (**Figure 4**). Among the 115 signals, the top association was between *STRN* and mean platelet thrombocyte volume (SBAT p-value = 5.9 × 10^−13^); previous studies have reported variant associations at this gene with the phenotype^19,20^.

**Figure 4.**
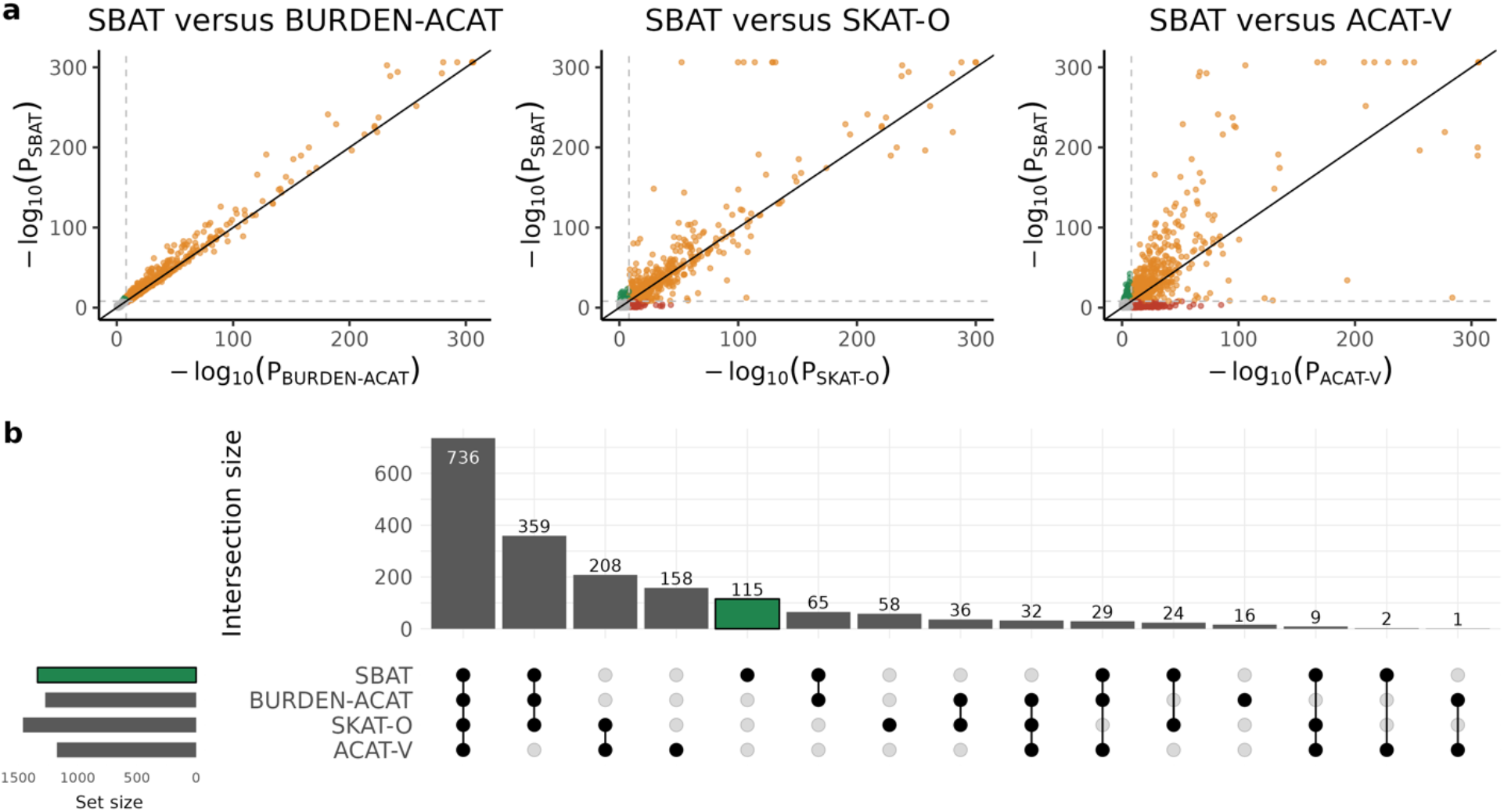
Gene-based analysis of 73 quantitative phenotypes with 18,184 genes using the UK Biobank data set. (a) Scatterplot of p-values comparing SBAT with BURDEN-ACAT, SKAT-O and ACAT-V. Each dot on the plot represents a gene result for a given trait where the y-axis denotes to the p-value (on -log10 scale) for SBAT and the x-axis denotes the p-value (on -log10 scale) for the comparison test. For BURDEN-ACAT, SKAT-O and ACAT-V, we applied ACAT to the p-values across the 4 annotation classes (and 5 MAF cutoffs for BURDEN-ACAT) to obtain a single p-value per gene. The dashed lines represent a significance threshold of 7.5 × 10^−9^ corresponding to a Bonferroni correction for about 5 million tests based on 73 traits analyzed, 18,184 genes and 4 methods applied. Green points refer to signals only found by SBAT, yellow points refer to signals detected by both tests and red points represent signals missed by SBAT but detected by the other test. P-values were capped at 2.2 × 10^−307^. (b) Upset plot of 1,848 genome-wide significant signals discovered across SBAT, BURDEN-ACAT, SKAT-O and ACAT-V association tests using a significance threshold of 9.*l* × 10^−9^.

We highlight two examples where using SBAT to jointly combine burden signals was the only method that led to a detectable signal compared to BURDEN-ACAT, SKAT-O, ACAT-V or the marginal burden tests using various MAF cutoffs and functional annotations. In **Figure 5**, the association between standing height^23,24^ and *PITX1*, which encodes a protein critical in the development of the lower limbs^21,22^, was only discovered by SBAT (p-value = 2.3 × 10^−11^). The strongest signal amongst the individual burden scores, which came from considering a burden of singleton pLoF variants, did not reach the genome-wide significance threshold (p-value = 3.8 × 10^−8^). Furthermore, both SKAT-O and ACAT-V gave p-values above that of the strongest burden signal (smallest p-values = 3.2 × 10^−7^ and 1.5 × 10^−7^, respectively). The joint model fitted with SBAT indicated that the gene signal was driven primarily by singleton pLoF variants as well as the combination of pLoF and missense variants predicted deleterious with a MAF below 0.1%, both associated with lower height. The second example was between platelet count and *ETV6*, which encodes a transcription factor^25^ (**Figure 6**). The association signal did not reach genome-wide significance threshold across the burden scores, where the smallest p-value was from a burden of pLoF and missense variants with a MAF cutoff of 1% (p-value = 2.6 × 10^−8^). Only by combining the burden scores signals though SBAT did the association reach the significance threshold (p-value = 1.3 × 10^−12^). Indeed, combining the burden scores p-values using the Cauchy combination method ACAT (BURDEN-ACAT) was not sufficient to reach the significance threshold (p-value = 2.7 × 10^−7^). Effect size estimates from the SBAT joint model indicated the signal was driven by singleton variants as well as more common pLoF and missense variants predicted deleterious.

**Figure 5.**
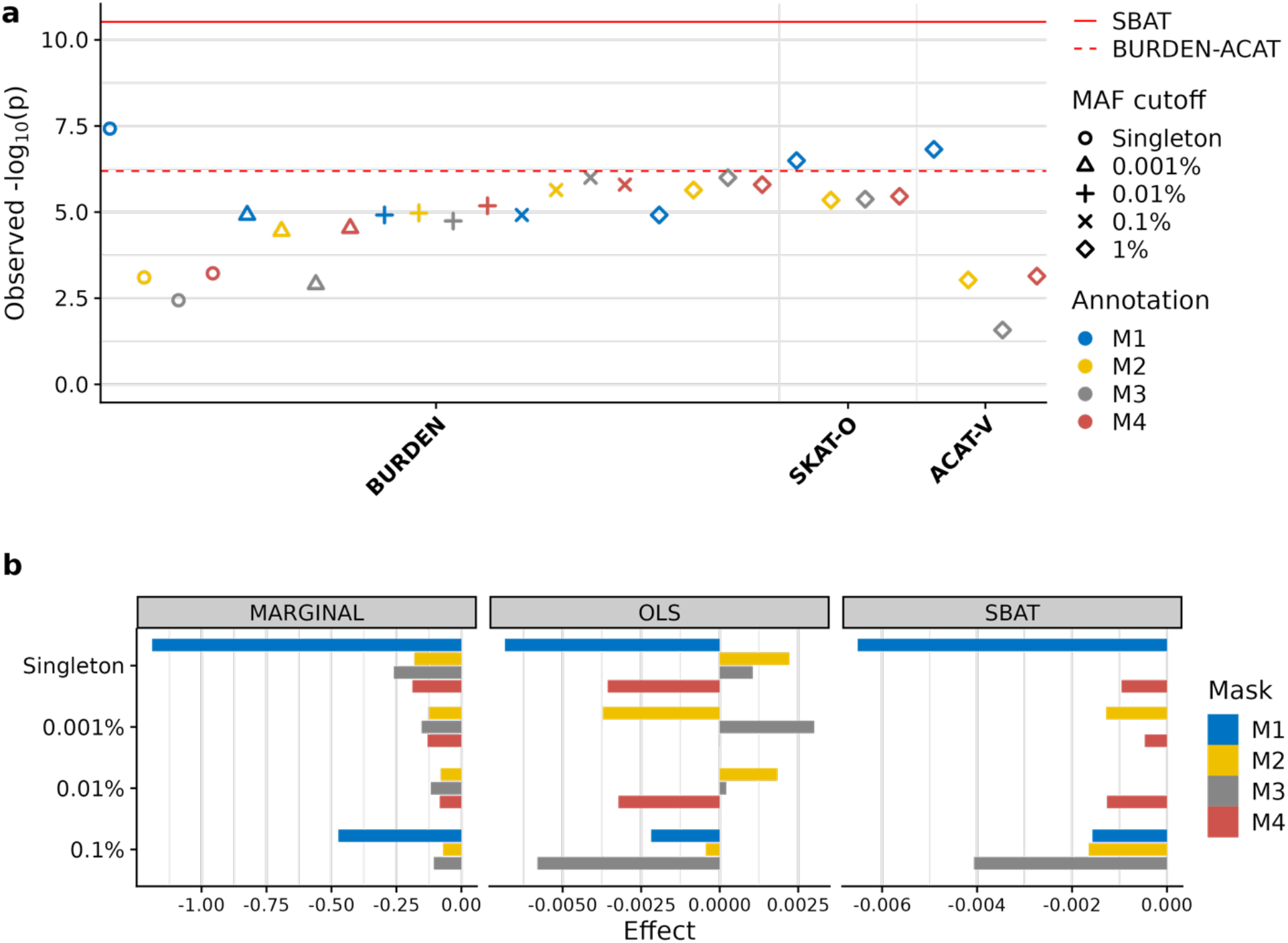
Deep dive into the association between *PITX1* and standing height. (a) Summary of all gene-based tests performed for the gene grouping variants based on MAF as well as annotation class (M1: pLoF only; M2: pLoF or missense; M3: pLoF or predicted deleterious missense based on 5/5 in-silico algorithms; M4: pLoF or predicted deleterious missense based on 1 or more out of 5 in-silico algorithms) using SKAT-O, ACAT-V, Burden test from REGENIE, BURDEN-ACAT, and SBAT on the burden scores. (b) Effect size estimates for the burden scores grouped by MAF and annotation class based on a marginal model with each mask (MARGINAL), a joint model on all scores using ordinary least squares (OLS), or the SBAT joint model (SBAT).

**Figure 6.**
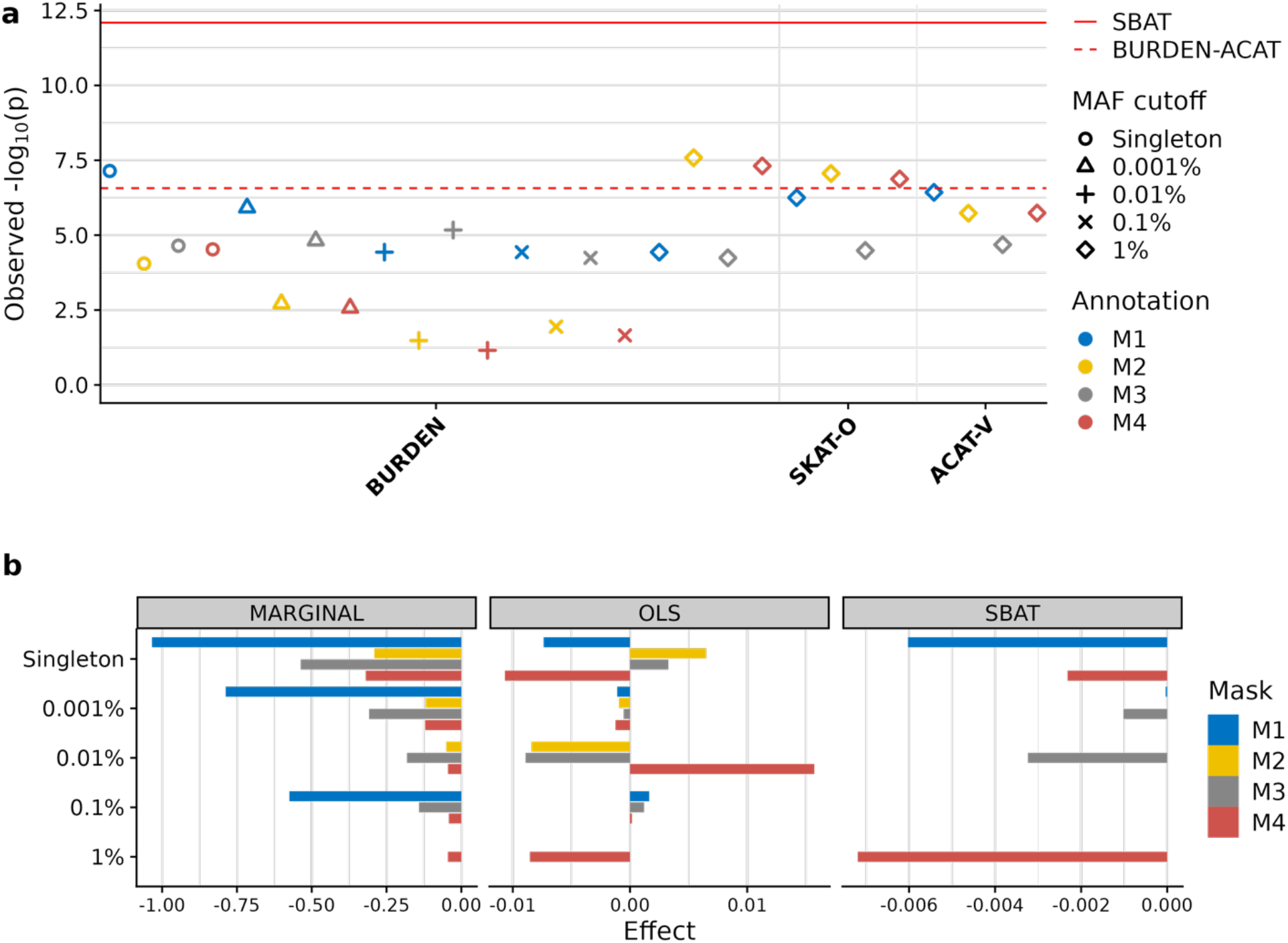
Deep dive into the association between *ETV6* and platelet count. (a) Summary of all gene-based tests performed for the gene grouping variants based on MAF as well as annotation class (M1: pLoF only; M2: pLoF or missense; M3: pLoF or predicted deleterious missense based on 5/5 in-silico algorithms; M4: pLoF or predicted deleterious missense based on 1 or more out of 5 in-silico algorithms) using SKAT-O, ACAT-V, Burden test from REGENIE, BURDEN-ACAT and SBAT on the burden scores. (b) Effect size estimates for the burden scores grouped by MAF and annotation class based on a marginal model with each mask (MARGINAL), a joint model on all scores using ordinary least squares (OLS), or the SBAT joint model (SBAT).

We also evaluated the performance of the unified test strategy, GENE_P, which combines results across SBAT, SKAT-O, ACAT-V and BURDEN-ACAT using the Cauchy combination method. GENE_P finds the most signals (1,685) compared to the other methods (1,313 for SBAT, 1,447 for SKAT-O, 1,156 for ACAT-V and 1,258 for BURDEN-ACAT) using a genome-wide significance threshold of 7.5 × 10^−9^ (**Supplementary Figure S4**). While there are 130 gene/trait associations that are missed by GENE_P, they all correspond to signals near the significance threshold (less than one order of magnitude away).

## Discussion

In this work, we introduced a new gene-based association test that pools information across multiple burden scores using constrained linear regression. Our approach is well-suited for gene-based testing: modeling burden scores jointly accounts for burden score correlations, while non-negative constraints induce sparsity in effect estimates. In simulations we showed that our SBAT method has well controlled type I error rates and outperforms SKAT-O and ACAT-V in scenarios where most causal variants have the same effect directions and rarer variants substantially contribute to the gene signal. We further confirmed our simulation results in the analysis of 73 quantitative phenotypes in the UK Biobank whole-exome sequencing data and showed that the four examined tests (SBAT, SKAT-O, ACAT-V and BURDEN-ACAT) share most of the gene signals, complementing each other by signals that are test-specific. SBAT notably revealed the highest number of unique associations (115) among the other two tests targeting the burden signals, SKAT-O (58) and BURDEN-ACAT (16). Following a common practice in designing gene-burden tests^9,10,11^, we proposed a strategy to provide a single p-value per gene by combining results from the four tests available at different variant annotation classes and allele frequency bins.

We highlighted two gene-phenotype SBAT associations in the UK Biobank analysis that were missed by alternative tests, SKAT-O, ACAT-V and BURDEN-ACAT. The two associations *PITX1*-standing height and *ETV6*-platelet count are well established in the literature^23,24,25^. Our results stress the role of pLoF singletons for both gene-phenotype pairs, where the burden scores with these variants show the lowest p-values (3.8 × 10^−8^ and 2.6 × 10^−8^, respectively) but do not pass the significance threshold of our study (9.4l × 10^−9^). The SBAT joint model aggregates the evidence of association from multiple burden scores amplifying the signal (SBAT p-values 2.3 × 10^−11^ and 1.3 × 10^−12^, respectively). These two SBAT findings agree with our simulation results, where SBAT outperforms other tests when the signal is concentrated at rarer variants with the same direction of effects. We also observe that SBAT gives more informative effect estimates in comparison to marginal and joint OLS estimates (**Figures 5b and 6b**), selecting the top burden scores of pLoF singleton variants and favoring missense burden scores with more likely deleterious annotations (M3 and M4 are preferred to M2).

Our study also has limitations. We used a particular set of nested annotations for burden scores from our previous works^2,3,12^ and considered only two main methods for comparison, SKAT-O and ACAT-V. While the list of selected annotations and methods is not exhaustive, we believe our simulation results served well to contrast the SBAT features against SKAT-O and ACAT-V. As data generation models can favor one method over the other in simulations, we empirically showed the competitive performance of SBAT in the exome-wide association study of 73 agnostically selected traits from 430,074 participants in the UK Biobank. The current implementation of SBAT is developed for quantitative traits, as the previous works on constrained linear regression were built for continuous response variable^15–17^. Future work will be focused on SBAT extension to binary traits, addressing issues such as testing rare variants for unbalanced binary traits^13^.

We envision that the constrained linear regression approach can also be applied to other association tests of multiple variables: joint testing of multiple traits^26^ and/or multiple SNPs ^27^. To enable these future extensions, we made an important practical contribution with approximate weight calculation in a mixture of chi-squared distributions for constrained linear regression, a problem not addressed in the previous theoretical works^15–17^. The constrained test can also operate using association summary statistics and correlation matrix of tested variables rather than individual-level data. We derived this summary-statistic option through reformulation of the constrained regression problem (Equation 1) in terms of joint effect sizes and the residual variance. The (unconstrained) joint effect estimates can be inferred from the marginal effect estimates using existing methods^28^.

In summary, we developed a scalable multivariate version of the one-sided test with application to testing multiple burden scores. The SBAT method is available in the REGENIE software and together with other gene-based tests will empower further discovery from sequencing association studies.

## URLs

REGENIE software https://rgcgithub.github.io/regenie/

---

**Supplementary Figure 1.**
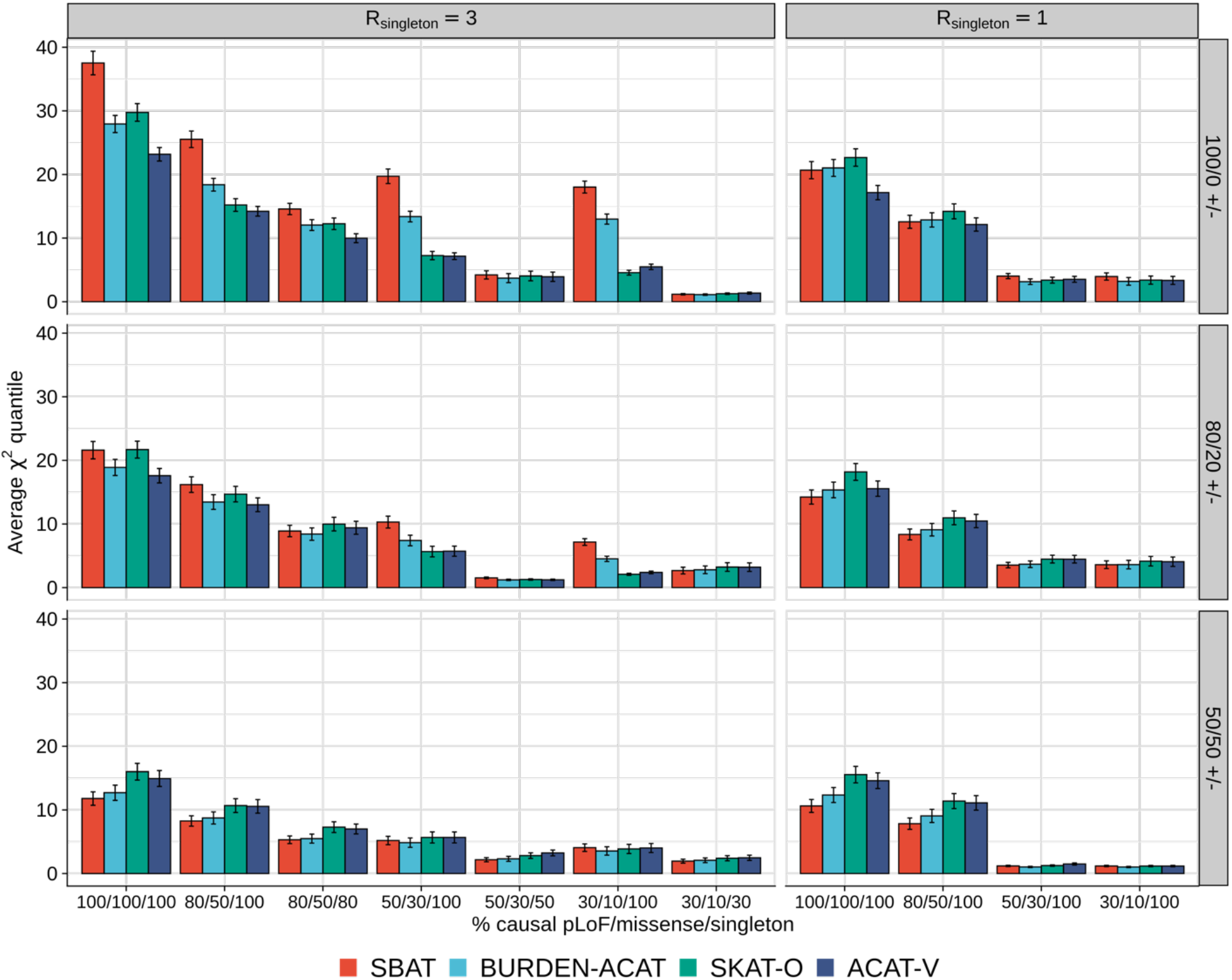
Power performance of SBAT in simulation studies under various causal scenarios. Average *χ*^2^ quantile for the set-based tests as a function of the proportion of causal variants amongst singletons, pLoF and missense variants. Each row corresponds to different assumptions for the direction of effects whether they are all positive (100/0 +/-), 80% positive and 20% negative (80/20 +/-), or half positive and the remaining negative (50/50 +/-). Each column corresponds to different assumptions for the effect of singleton variants, which was multiplied by a constant factor *R*_*singleton*_ relative to non-singletons. The error bars represent 95% confidence intervals for the average *χ*^2^ quantile. SBAT is compared to SKAT-O, ACAT-V and BURDEN-ACAT association tests, where BURDEN-ACAT applies the Cauchy combination test ACAT to the individual burden mask signals.

**Supplementary Figure 2.**
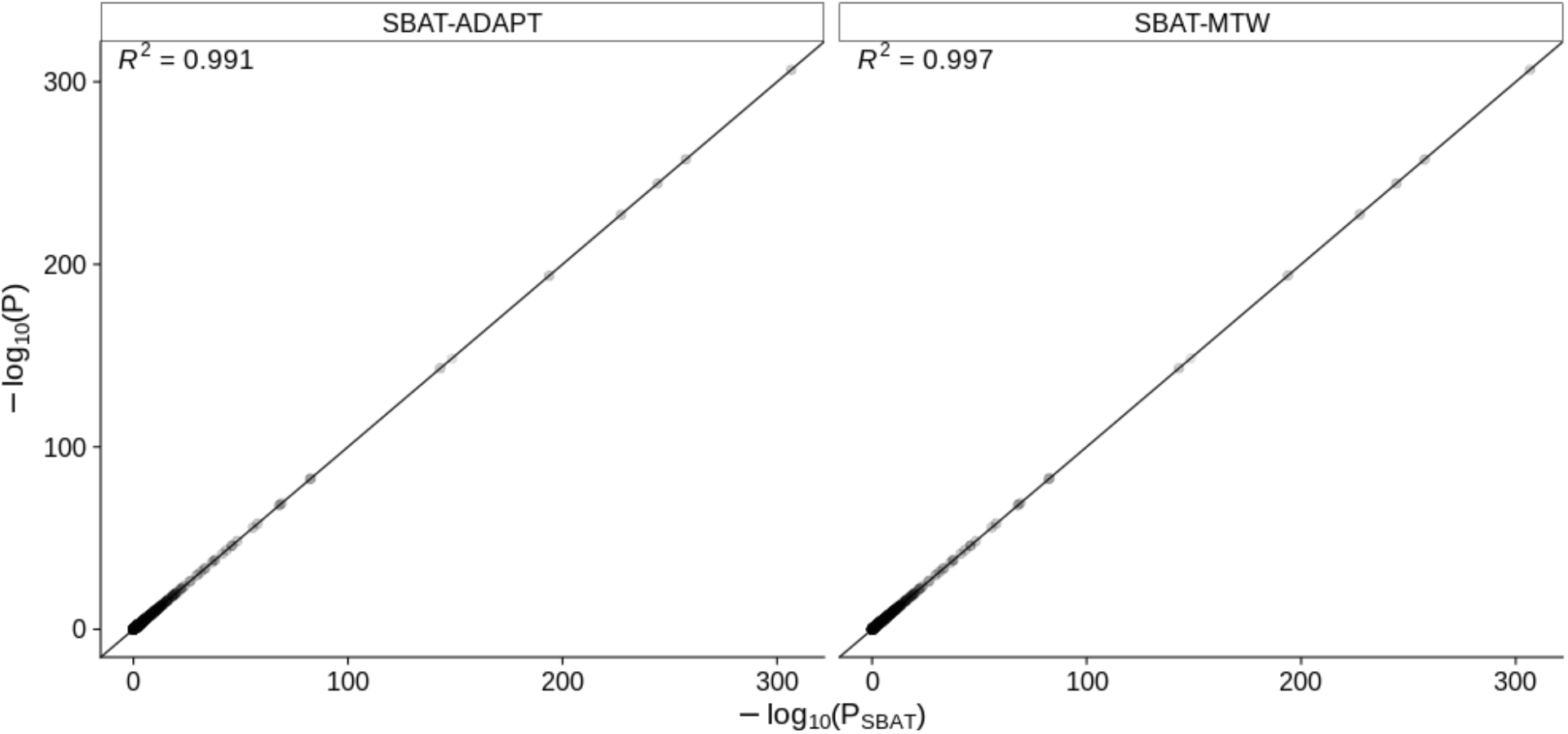
Gene-based association test results for 50 quantitative phenotypes in UK Biobank. The -log_10_ p-values obtained from SBAT (x-axis) are compared against those using either the adaptive strategy (SBAT-ADAPT) or the multi-trait weights strategy (SBAT-MTW) on the y-axis. Each point represents a gene where a joint test is applied on a set of 24 burden scores (from 4 annotation classes and 6 AAF cutoffs) across 804 genes on chromosome 16.

**Supplementary Figure 3.**
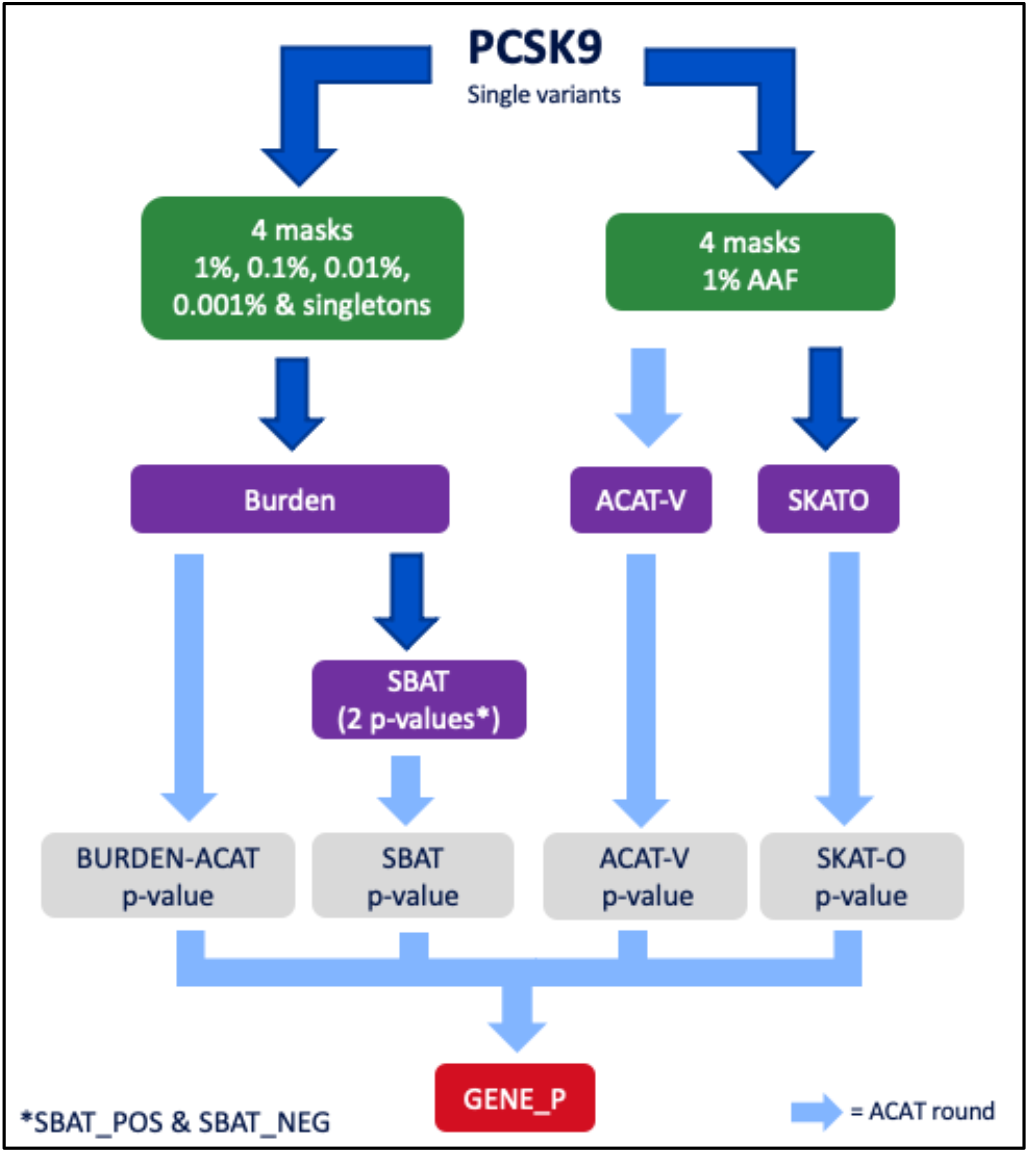
Unified strategy GENE_P to obtain a single p-value per gene across various gene-based tests. The Cauchy combination method ACAT as well as SBAT are applied to a set of burden scores, and ACAT-V and SKAT-O are applied to single variant results. Variant are grouped into four annotation classes and 5 MAF cutoffs (single cutoff is used for ACAT-V and SKAT-O).

**Supplementary Figure 4.**
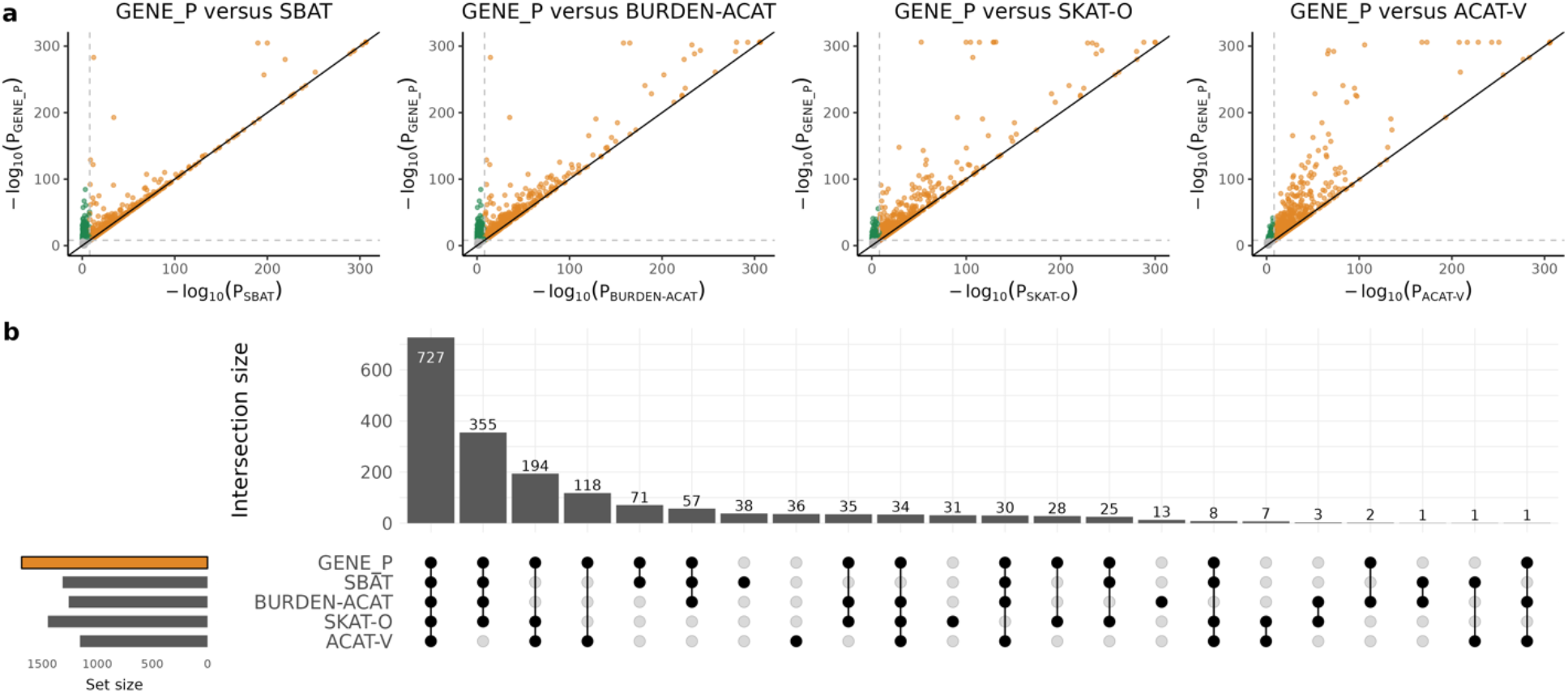
Gene-based analysis of 73 quantitative phenotypes with 18,184 genes using the UK Biobank data set. (a) Scatterplot of p-values comparing unified strategy GENE_P with SBAT, BURDEN-ACAT, SKAT-O and ACAT-V. Each dot on the plot represents a gene result for a given trait where the y-axis denotes to the p-value (on -log10 scale) for GENE_P and the x-axis denotes the p-value (on -log10 scale) for the comparison test. For BURDEN-ACAT, SKAT-O and ACAT-V, we applied ACAT to the p-values across the 4 annotation classes (and 5 MAF cutoffs for BURDEN-ACAT) to obtain a single p-value per gene. The dashed lines represent a significance threshold of 7.5 × 10^−9^ corresponding to a Bonferroni correction for about 6.6 million tests based on 73 traits analyzed, 18,184 genes and 5 methods applied. Green points refer to signals only found by GENE_P, yellow points refer to signals detected by both tests and red points represent signals missed by GENE_P but detected by the other test. P-values were capped at 2.2 × 10^−307^. (b) Upset plot of 1,815 genome-wide significant signals discovered across GENE_P, SBAT, BURDEN-ACAT, SKAT-O and ACAT-V association tests using a significance threshold of 7.5 × 10^−9^.

## Notes

### Competing Interest Statement

All authors are current or former employees and/or stockholders of Regeneron Genetics Center or Regeneron Pharmaceuticals.

### Summary of Updates

Supplementary Tables file added

